# Towards routine employment of computational tools for antimicrobial resistance determination via high-throughput sequencing

**DOI:** 10.1101/2021.11.03.467126

**Authors:** Simone Marini, Rodrigo A. Mora, Christina Boucher, Noelle Noyes, Mattia Prosperi

## Abstract

Antimicrobial resistance (AMR) is a growing threat to public health and farming at large. In clinical and veterinary practice, timely characterization of the antibiotic susceptibility profile of bacterial infections is a crucial step in optimizing treatment. High-throughput sequencing is a promising option for clinical point-of-care and ecological surveillance, opening the opportunity to develop genotyping-based AMR determination as a possibly faster alternative to phenotypic testing. In the present work, we compare the performance of state-of-the-art methods for detection of AMR using high-throughput sequencing data from clinical settings. We consider five computational approaches based on alignment (AMRPlusPlus), deep learning (DeepARG), *k*-mer genomic signatures (KARGA, ResFinder) or hidden Markov models (Meta-MARC). We use an extensive collection of 585 isolates with available AMR resistance profiles determined by phenotypic tests across nine antibiotic classes. We show how the prediction landscape of AMR classifiers is highly heterogeneous, with balanced accuracy varying from 0.40 to 0.92. Although some algorithms—ResFinder, KARGA, and AMRPlusPlus– exhibit overall better balanced accuracy than others, the high per-AMR-class variance and related findings suggest that: (1) all algorithms might be subject to sampling bias present both in data repositories used for training and experimental/clinical settings; and (2) a portion of clinical samples might contain uncharacterized AMR genes that the algorithms—mostly trained on known AMR genes—fail to generalize upon. These results lead us to formulate practical advice for software configuration and application, and give suggestions for future study designs to further develop AMR prediction tools from proof-of-concept to bedside.

## Introduction

Antimicrobial resistance (AMR) occurs when microorganisms evolve to overcome susceptibility to antibiotics. According to the US Centers for Disease Control (CDC), more than 2.8 million antibiotic-resistant infections and over 35,000 resultant deaths occur each year in the US alone[1]. On a national level, it is predicted that AMR-related infections will lead to 10 million deaths per year and a gross domestic product (GDP) loss of $100.2 trillion by 2050 without appropriate interventions [2]. From managing antibiotic misuse in healthcare to regulating their use in livestock and agriculture, AMR has become an “arms race” between microorganism adaptation and drug discovery. In clinical settings, one major obstacle to optimizing treatment of AMR-related infections is access to accurate and timely antibiotic susceptibility testing (AST)[3]. Wet-lab AST techniques generally require growing bacteria in vitro and testing against various antibiotics. Examples include agar-based culture techniques, disk diffusion, and Etest [4]. These techniques are limited in that: (a) culture conditions have not been validated for every infectious microbial species, and only a fraction of bacterial species are cultivable with standard methods; (b) they may take as long as 5 days before the result is obtained (1–3 days for culture and 1-2 days for AST)[3]; (c) result interpretation standards are constantly evolving; and (d) they are resource intensive in terms of training personnel effort, equipment, and consumables.

The widespread use of DNA sequencing technologies has led to the development of curated databases of AMR genes complementing culture experiments, with detailed functional annotation and other metadata[5, 6, 7, 8, 9, 10]. In turn, computational approaches have emerged to provide in silico detection and characterization of AMR phenotypes and genes, processing targeted sequences as well as metagenomic samples from both clinical and environmental settings. These tools can directly process high-throughput short read sequencing data (typically in FASTQ format)[9, 11, 12, 13], or take as input assembled contigs, translated proteins, or whole bacterial genomes (typically FASTA format)[8, 14, 15, 16, 17, 18]. Also, some tools are purposely designed for metagenomics data, while others specialize on specific bacterial species. Despite the promising results shown in literature and support from experimental studies, the uptake of sequencing-based computational AMR prediction tools for routine use in healthcare has been slow[19]. One of the major roadblocks is that algorithms can exhibit discordant performance in clinical settings, and there is not yet a gold standard or consensus like the Clinical & Laboratory Standards Institute (CLSI) for AST. Sub-problems that impede the development of such gold standard include: (a) possible shifts and bias in species or genus representations when comparing the training settings of models with the application settings[10, 20]; (b) different working assumptions of the tools and required inputs; and (c) choice of biological ontologies for AMR representation, i.e., standardized encoding of AMR agents and mechanisms, and prediction outputs. These problems make it difficult to compare methods in terms of performance, agreement, and generalizability, complicating the establishment of benchmark data sets and external validation. Prior works identified strong spatiotemporal heterogeneity in sampling and species representation in genotype-phenotype data repositories, confounding the identification of AMR gene signatures, as well as the generalizability of prediction models on independent data[20, 21]. For example, some AMR genes or variants are species-specific, e.g., resistance to fluoroquinolones via the *gyrA* gene is specific to *Mycoplasma genitalium*[22, 9, 15].

From one perspective, AMR detection algorithms operating on whole genomes and using rules for specific bacterial strains have a design advantage with respect to species representation bias. However, requiring a whole genome sequence as input implies that the bacterial strain has been identified prior to sequencing, which excludes unculturable pathogens. Furthermore, software for genome assembly must be run, either de novo or reference-based, with potential downstream errors.

Algorithms which take as input unassembled sequence reads do not need a priori information about the bacterial species in the sample. In addition, metagenomics-based, culture-independent genomic approaches can be used concurrently with the traditional methods to help detect AMR in unculturable bacteria, eliminating the necessity of species isolation [23]. The absence of assembled genomes makes the AMR classification task more challenging, since results must be aggregated from the read to the isolate level, and species-specific AMR rules might need to be embedded in the algorithms. Nonetheless, these algorithms are in principle more flexible and applicable to a variety of clinical scenarios, involving both characterization of known AMR as well as possibly novel microbial elements or resistance genes.

Regardless of the methodology of operation, all methods are subject to ontology issues, as there are multiple ontologies for AMR genes and mechanisms, and some tools predict AST phenotypes directly rather than AMR genes.

In this study, we assess the predictive performance of state-of-the-art AMR classification algorithms for high-throughput bacterial sequencing data from clinical settings. We consider different tools that include alignment-based (AMRPlusPlus)[9], *k*-mer signatures (KARGA, ResFinder)[13, 24], and machine learning (Meta-MARC, DeepARG)[11, 12] methods. We have designed an extensive benchmark setup apt to address agreement and generalizability issues. More specifically, we have collated high-throughput sequencing data from 585 clinical isolates, sourced from different studies in clinical settings worldwide (Asia, Europe, North America, and South America), all with available AMR resistance profiles determined by phenotypic AST, covering nine major AMR classes. We provide a robust assessment of the algorithms capabilities and discuss the potential future implications for their routine use in healthcare settings.

## 1 Methods

### 1.1 AMR algorithms and software

Notwithstanding a very large collection of AMR prediction methods, this study includes approaches that directly take high-throughput sequencing data as input, removing the necessity for genome assembly prior to analysis. We compare the following AMR computational tools: AMRPlusPlus[9], DeepARG[12], KARGA[13], Meta-MARC[11], and ResFinder[24]. As previously mentioned, these methods use a variety of different computational approaches, i.e. alignment-based, *k*-mer based (*k*-mers are nucleotide signatures of fixed length k), and machine learning. Also, some tools provide predictions at the read level (DeepARG), while others at the isolate level (AMRPlusPlus, ResFinder), or both (KARGA, Meta-MARC). We compare the methods in Table 1.

**Table 1:** Summary of AMR prediction tools evaluated in this study, describing: publication year; algorithmic approach; reference AMR database; output capabilities as whole-sample and/or per-read AMR gene prediction; working platform(s). *HMM*, Hidden Markov models; *CNN* (Convolutional Neural Network; *JVM Java Virtual Machine.*

| Tool<br>(Publication year) | Algorithmic approach | AMR<br>database | Whole-sample<br>resistome<br>characterization | Per-read<br>resistome<br>classification | Platform |
| --- | --- | --- | --- | --- | --- |
| **AMRPlusPlus** (2020) | Read alignment | MEGARes | ✓ |  | UNIX/Linux |
| **Meta-MARC** (2019) | HMM classification | MEGARes | ✓ | ✓ | UNIX/Linux |
| **KARGA** (2021) | Comparison of $k$-mers | MEGARes,<br>or any other<br>fasta AMR DB | ✓ | ✓ | Any with JVM |
| **DeepARG** (2018) | Read alignment +<br>CNN classification | DeepARG DB |  | ✓ | Any with pip,<br>Conda, Docker |
| **ResFinder** (2020) | $k$-mers + alignment | ResFinder DB +<br>PointFinder DB | ✓ | | Any with Python 3,<br>Docker; Webserver |

**AMRPlusPlus (v.2.0)[9]** is a comprehensive software framework built around the MEGARes database[9]. MEGARes includes ~8,000 AMR genes and variants organized in a multi-level hierarchical ontology in the form of a directed acyclic graph, comprising four compound types, 57 classes, 220 mechanisms, and 1,345 groups. This structure ensures that two higher level ranks are not linked to the same lower level rank, and univocity in sequence classification. AMRPlusPlus aligns all reads to MEGARes using the BWA[25], retaining AMR genes with a coverage of at least 80%. Low-quality and low-depth alignments are filtered through the Resistome Analyzer and the Rarefaction Analysis modules, respectively.

**Meta-MARC[11]** is a machine-learning model based on an ensemble of hierarchical hidden Markov models (HMMs). Each HMM is trained on a group of genes clustered from MEGARes[11]. A classification is performed by aggregating predictions from the lowest level of the MEGARes annotation hierarchy towards the highest level, although there is not always a complete match with the MEGARes ontology due to gene aggregation/split. Meta-MARC provides a three-model ensemble (model groups I-III). Group I models are based on AMR sequences clusters containing more than two unique sequences for each single type of AMR; Group II expands group I and includes genes carrying AMR variants due to single nucleotide polymorphisms. Group III includes all of groups I and II, plus additional sequences obtained by aligning MEGARes genes against the National Center for Biotechnology Information (NCBI) GenBank using the Basic Local Alignment Search Tool (BLAST) and retaining highly similar sequences.

**KARGA[13]** is an alignment-free tool. It uses sequenced gene *k*-mers, storing the whole *k*-mer spectrum—i.e., all distinct *k*-mers—both in forward- and reverse-strand from any given reference AMR database[13]. To classify a new sequence, KARGA calculates its *k*-mer spectrum and compares it with the AMR database spectrum. To avoid false positives, KARGA calculates an empirical distribution of matches between randomly generated *k*-mers and the AMR database, approximating a theoretical Markovian distribution, finding an optimal threshold based on query length, *k* value, and the database spectrum. A probabilistic scoring manages multiple gene hits. For this study we used MEGARes, which was also employed in the original KARGA publication.

**DeepARG[12]** is a two-step method that combines read alignment and convolutional deep learning networks. In the first step, an input sequence is translated to amino acids, then aligned to a custom AMR database, which was created by merging the Comprehensive Antibiotic Resistance Database (CARD)[5, 12], Antibiotic Resistance Genes Database (ARDB)[6], and manually curated AMR sequences from the Universal Protein Resource (UNIPROT). If the sequence aligns to the custom database, it is passed on to the second step where the deep learning module predicts the AMR category. Of note, the DeepARG classification output does not follow either MEGARes or CARD ontology, but it is a customized set of thirty AMR categories. DeepARG features two working modes, one for long reads and one for short reads.

**ResFinder (v.4.0)[24]** is a *k*-mer based approach based on Python 3 scripts, that can process both sequence reads and assembled contigs/genomes. The Resfinder *k*-mer engine is based on KMA[24, 26]. Briefly, KMA implements a heuristic *k*-mer matching score with a threshold to accept query sequences for subsequent alignment. Seeds of the identified *k*-mers are used to guide a modified Needleman-Wunsch alignment, which includes a score-based early stop. An additional scoring system, ConClave [26], is then used to resolve tied alignments on redundant sequences and assemble the final output. Notably, bacterial species can be specified by the user in the input, thus customizing the output results for a subset of species-specific resistances.

### 1.2 Study selection, data set collation, quality control, and evaluation

We performed a literature search for AMR sequencing studies on PubMed Central based on keywords and MESH terms related to AMR (see supplementary material for complete list). We then selected studies with the goal of collating a data set of at least 500 isolates according to the following criteria: AMR-focused; clinical setting; published within the last seven years (2014-2021); sequenced using Illumina platform; available NCBI BioProject and/or Accession number/code identifier; at least 25 paired-end experiments available in NCBI Sequence Read Archive (SRA); and available AMR resistance profiles determined by phenotypic AST. All FASTQ files were processed with Trimmomatic v.0.32, using all Illumina adapters, and the following parameters: LEADING=3, TRAILING=3 SLIDINGWINDOW=4:15, MINLEN=36. Unpaired reads were discarded. We considered isolates as resistant if they were marked resistant (R), intermediate (I) or susceptible dose dependent (SDD), and sensitive if they were marked sensitive (S) in their source paper. An isolate was considered resistant or susceptible to an antibiotic class if it was marked in its source paper as resistant or sensitive, respectively to at least one antibiotic within that antibiotic class.

We used the CARD and MEGARes ontologies to link antibiotics to their AMR classes, and considered only the classes that could be predicted by all the tools included in this study. Consistency adjustments of AMR labeling among different algorithms due to different ontology terms were curated by hand (for example, the quinolone DeepARG category corresponds to the fluoroquinolone class used by the other methods). Each isolate was labeled as resistant or susceptible to specific AMR classes. Therefore, we could label isolates as positive (resistant) and negative (susceptible) with respect to each AMR class. To evaluate the performance of the methods, we considered Sensitivity (Sens, or true positive rate), Specificity (Spec, or true negative rate), Balanced Accuracy (BalAcc, the mean of Sens and Spec), and *f*_1_ score, which is the harmonic mean between precision (positive predicted value) and recall (Sens).

### 1.3 Software and parameter setup

AMRPlusPlus was run with its default parameters. DeepARG was run in short read mode, using the recommended threshold probability of (0.80 for read classification, and the rest of the parameters were set to default. Meta-MARC was run specifying paired end input, and model group III, with other parameters set to default. KARGA was run with its default parameters (*k* had been previously optimized), filtering AMR genes with coverage below 80% as recommended. ResFinder was used with default parameters.

## 2 Results

### 2.1 Data set characteristics, spatio-temporal and species distributions

Five-hundred-eighty-five isolates from six sources (here labeled as S1-S6) met all the study selection criteria. In summary, Weingarten et al. (S1) performed whole genome sequencing on 108 clinical isolates containing carbapenemase-producing organisms (CPOs) from a clinical center in Bethesda, Maryland, USA [27]. Runcharoen et al. (S2) analyzed the relationship between clinical and environmental *Klebsiella pneumoniae* strains, performing whole genome sequencing and phylogenetic analysis of 77 isolates sampled from a hospital, its sewers, and the surrounding canals and farms within a 20-km radius in Thailand [28]; we considered only hospital-related isolates in our analysis. Davies et al. (S3) performed genome sequencing and phylogenetic analysis of 141 clinical isolates from patients, 58 presenting with scarlet fever and 83 with other clinical presentations from Hong Kong, Australia, United States, and Mainland China [29]. Croucher et al. (S4) performed whole genome sequencing and phylogenetic analysis of 189 clinical isolates from twelve countries from 1988 to 2009, discussing the potential relationship between antibiotic dispensing prevalence and antibiotic resistance profiles [30]. Alonos-del Valle et al. (S5) analyzed the antibiotic resistance profiles of 50 gut isolates from patients in a hospital in Madrid, Spain, comparing sample-originated resistance profiles to those produced by introducing a carbapenem-resistance plasmid into each enterobacterial isolate; we considered only sample-originated resistance profiles [31]. Pesesky et al. (S6) predicted antibiotic susceptibility of 78 de-identified patient samples from clinical bacterial biobanks in Rawalpindi and Islamabad, Pakistan and from Saint Louis, Missouri, USA. Enterobacteriaceae isolates, which had been previously sequenced, were analyzed using two genotypic computational algorithms and the results were compared to previously identified phenotypic AST profiles [16]. Additional details of the data sets, such as labeling and filtering used to assemble them for this study are provided in the Supplementary Methods and Supplementary Tables S1-2.

According to the AMR ontology linkage by MEGARes and CARD defined in the methods, and based on the phenotypic test determined resistance profiles of the study isolates, the following classes are retained in the analysis: aminoglycosides (amikacin, tobramycin, and gentamicin); betalactamases (amoxicilin/clavulanic acid, ampicillin, aztreonam, cefalotin, cefazolin, cefepime, cefotaxime, cefotetan, cefoxitin, ceftazidime, cef-triaxone, cefuroxime, cefuroxime/axetil, doripenem, ertapenem, imipenem, meropenem, penicillin, and piper-cillin/tazobactam); fluoroquinolones (ciprofloxacin and levofloxacin); glycopeptides (vancomycin); macrolide, lincosamide, and streptogramin –MLS– (clindamycin and erythromycin); phenicols (chloramphenicol); sulfonamides (trimethoprim/sulfamethoxazole); tetracyclines (doxycycline, tigecycline, and tetracycline); and diaminopyrimidines (trimethoprim and trimethoprim/sulfamethoxazole). We considered isolates resistant to trimethoprim/sulfamethoxazole as resistant to both sulfonamides and diaminopyrimidines (trimethoprim), while the susceptible ones were not considered, as the resistance to individual trimethoprim and sulfamethoxazole were not measured in the corresponding original data sets.

Tables 2 and 3 summarize the characteristics of each study and per-AMR-class isolates details, respectively. More details are available in the Supplementary Methods, and in Supplementary Tables S1-2.

**Table 2:** Summary of sources included in our AMR benchmarking study. *^a^*For isolate and data pre-processing details, see Supplementary Methods and Supplementary Table S1. *^b^*For isolate antibiotic resistance profiles per organism and antibiotic class, see Supplementary Table S3. *^c^*VITEK 2 (bioMérieux, Marcy l’Étoile, France). *^d^*Only isolates with available Resistant (R), Intermediate (I), and/or Sensitive (S) labels were considered for analysis.

|  | Isolates<sup>a</sup><br>BioProject(s)<br>(Publication Year) | Data obtained<br>from S1-S6 Publications | Data obtained from NCBI SRA Run Selector |  |
| --- | --- | --- | --- | --- |
|  |  | Phenotypic Antibiotic<br>Susceptibility Testing | Country | Organism (Isolates)<sup>b</sup> |
| **S1** | 142<br>PRJNA430442<br>(2018) | Broth microdilution<br>Disk diffusion | USA | Achromobacter sp. (2)<br>Acinetobacter baumannii (2)<br>Acinetobacter calcoaceticus/baumannii<br>complex sp. (1)<br>Acinetobacter sp. (4)<br>Aeromonas hydrophila (3)<br>Aeromonas sp. (10)<br>Citrobacter freundii (1)<br>Citrobacter freundii complex sp. (13)<br>Citrobacter sp. (2)<br>Enterobacter cloacae (1)<br>Enterobacter cloacae complex sp. (19)<br>Enterobacteriaceae bacterium (5)<br>Escherichia coli (8)<br>Escherichia sp. (1)<br>Klebsiella aerogenes (2)<br>Klebsiella oxytoca (15)<br>Klebsiella pneumoniae (23)<br>Klebsiella pneumoniae subsp.<br>Pneumoniae (2)<br>Leclercia sp. (11)<br>Pantoea sp. (5)<br>Pseudomonas sp. (2)<br>Serratia sp. (10) |
| **S2** | 50<br>PRJEB11403<br>(2017) | VITEK 2<sup>c</sup> | Thailand | Klebsiella pneumoniae (45)<br>Klebsiella quasipneumoniae (5) |
| **S3** | 133<br>PRJEB2657,<br>PRJEB2589<br>(2014) | Disc diffusion,<br>D-zone disc diffusion<br>Etest | Hong Kong | Streptococcus pyogenes (133) |
| **S4** | 140<br>PRJEB2255<br>(2014) | Phenotypic tests for<br>isolates labeled 'resistant',<br>'intermediate' or 'sensitive'<br>not specified<sup>d</sup> | Iceland<br>Thailand<br>Portugal<br>Peru | Streptococcus pneumoniae (140) |
| **S5** | 47<br>PRJNA641166<br>(2021) | Disc diffusion | Spain | Escherichia coli (23)<br>Klebsiella pneumoniae (19)<br>Klebsiella quasipneumoniae (4)<br>Klebsiella quasivariicola (1) |
| **S6** | 73<br>PRJNA261540<br>(2016) | Kirby Bauer<br>Disk Diffusion | Pakistan | Enterobacter cloacae (8)<br>Escherichia coli (33)<br>Klebsiella aerogenes (4)<br>Klebsiella pneumoniae (28) |

**Table 3:** Summary of the resistant and sensitive isolates, most represented country, and most prevalent genus per AMR class from our collated data set derived from S1-S6 data sets.

| Antibiotic Class | Resistant Isolates | Sensitive Isolates | Most Represented Country | Most Prevalent Genus |
| --- | --- | --- | --- | --- |
| Aminoglycosides | 181 (58%) | 131 (42%) | USA (53%) | Klebsiella (47%) |
| betalactamases | 326 (66%) | 166 (34%) | USA (33%) | Streptococcus (37%) |
| Fluoroquinolones | 146 (55%) | 119 (45%) | USA (62%) | Klebsiella (47%) |
| Glycopeptides | 0 (0%) | 133 (100%) | Hong Kong (100%) | Streptococcus (100%) |
| MLS | 217 (81%) | 51 (19%) | Hong Kong (50%) | Streptococcus (100%) |
| Phenicol | 199 (63%) | 119 (37%) | Iceland (36%) | Streptococcus (42%) |
| Sulfonamides | 141 (100%) | 0 (0%) | USA (72%) | Klebsiella (35%) |
| Tetracyclines | 301 (77%) | 88 (23%) | Hong Kong (34%) | Streptococcus (68%) |
| Diaminopyrimidines (Trimethoprim) | 180 (94%) | 11 (6%) | USA (53%) | Klebsiella (52%) |

The data sets present highly heterogeneous per-class geographical, temporal, and bacterial distributions. To assess distribution shifts and putative bias, we performed tests for equal proportions of AST resistance by country of origin, year of collection, and genus. Results indicate an extreme heterogeneity for at least one of the three domains for each AMR class, with the exclusion of diaminopyrimidines (trimethoprim). More specifically, stratified resistance prevalence exhibited wide ranges for the following AMR classes: aminoglycosides (country [0.14, 0.72], year [0, 0.85], and genus [0, 1]); betalactamases (country [0, 1], year [0, 1], and genus [0.25, 1]); fluoroquinolones (year [0.28, 1] and genus [0, 1]); MLS (year [0, 1]); phenicols (country [0.23, 0.87], year [0, 1], and genus [0, 0.89]); and tetracyclines (year [0, 1]). Nonetheless, the odds-ratio (OR) ranges for AMR prevalence shift across calendar years did not indicate a large effect size, varying between 0.98 and 1.40, with five out of seven ORs being larger than 1, suggesting a slight shift towards increased resistance per more recent calendar year (only seven AMR classes considered in this study have both resistant and susceptible isolates.)

### 2.2 Computational determination of AMR, prediction performance, and agreement among algorithms

All 585 isolates were processed by each AMR computational tool. The prediction performances are summarized in Table 4. Overall, ResFinder yields the highest average BalAcc across all classes, 0.87 (InterQuartile Range, IQR 0.63, 0.89), followed by KARGA with 0.83 (0.73, 0.84), AMRPlusPlus with 0.80 (0.51, 0.89), DeepARG with 0.55 (0.50, 0.62), and Meta-MARC with 0.50 (0.50, 0.62). Comparing the per-AMR-class BalAcc in the top three methods, the null hypothesis that one distribution is greater than the other cannot be rejected (Wilcoxon rank sum test p-value of 0.74 and 0.34 for ResFinder versus KARGA and ResFinder versus AMRPlusPlus, respectively). We observe a wide per-AMR-class heterogeneity, without any method performing superior to the others on all the classes. KARGA is the most robust according to quartile coefficient of dispersion (0.07 for KARGA, 0.17 for ResFinder). For aminoglycosides and fluoroquinolones, all algorithms perform poorly, with the best BalAcc achieved by KARGA (0.66 and 0.58, respectively) and ResFinder (0.64 and 0.61, respectively). For betalactamases, the best algorithm is AMRPlusPlus with a BalAcc of 0.90; other algorithms perform in the [0.83, 0.89] range, with the exception of ResFinder (0.50). For MLS, ResFinder and KARGA achieve a BalAcc of 0.93 and 0.83, respectively, while other algorithms are limited to the [0.40, 0.57] range. For phenicols, tetracyclines, and diaminopyrimidines (trimethoprim), ResFinder, KARGA, and AMRPlusPlus achieve BalAccs in the [0.80, 0.91] range, while other algorithms are limited to the [0.50, 0.72] range. Glycopeptides, where our data set consists of only susceptible isolates, and sulfonamides, where our data set consists of only resistant isolates, were not considered in the main analysis. For completeness, we report the performances of these antibiotic classes in the Supplementary Methods and Supplementary Table S4.

**Table 4:** Classification performance for all methods on S1-S6 data sets (585 isolates), stratified per AMR class. Instances in bold show the top-performing methods whose balanced accuracy overlaps according to the standard error (SE).Considering True Positives as TP, True Negatives as TN, False Positives as FP, False Negatives as FN, we define Sensitivity (Sens) as 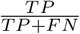; Specificity (Spec) as 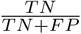; F-score as 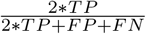; and Balanced Accuracy (BalAcc) as 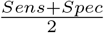

| Antibiotic Class | Algorithm | Sens ± SE | Spec ± SE | F-score ± SE | BalAcc ± SE |
| --- | --- | --- | --- | --- | --- |
| Aminoglycosides | AMRPlusPlus | 0.99 ± <0.01 | 0 | 0.64 ± 0.03 | 0.50 ± 0.03 |
| | DeepARG | 1 | 0 | 0.64 ± 0.03 | 0.50 ± 0.03 |
| | Meta-MARC | 0.99 ± 0.01 | 0.02 ± 0.01 | 0.64 ± 0.03 | 0.50 ± 0.03 |
| | **KARGA** | 0.98 ± 0.01 | 0.35 ± 0.03 | 0.73 ± 0.03 | **0.66 ± 0.03** |
| | **ResFinder** | 0.99 ± 0.01 | 0.30 ± 0.03 | 0.71 ± 0.03 | **0.64 ± 0.03** |
| Betalactamases | **AMRPlusPlus** | 1 | 0.80 ± 0.02 | 0.95 ± 0.01 | **0.90 ± 0.01** |
| | DeepARG | 0.87 ± 0.01 | 0.80 ± 0.02 | 0.89 ± 0.01 | 0.84 ± 0.02 |
| | **Meta-MARC** | 0.98 ± 0.01 | 0.80 ± 0.02 | 0.94 ± 0.01 | **0.89 ± 0.01** |
| | KARGA | 0.86 ± 0.02 | 0.81 ± 0.02 | 0.88 ± 0.01 | 0.83 ± 0.02 |
| | ResFinder | 1 | 0 | 0.80 ± 0.02 | 0.50 ± 0.02 |
| Fluoroquinolones | AMRPlusPlus | 0.97 ± 0.01 | 0.08 ± 0.02 | 0.72 ± 0.03 | 0.53 ± 0.03 |
| | DeepARG | 1 | 0 | 0.71 ± 0.03 | 0.50 ± 0.03 |
| | Meta-MARC | 0.98 ± 0.01 | 0.02 ± 0.01 | 0.70 ± 0.03 | 0.50 ± 0.03 |
| | **KARGA** | 0.60 ± 0.03 | 0.55 ± 0.03 | 0.61 ± 0.03 | **0.58 ± 0.03** |
| | **ResFinder** | 0.92 ± 0.02 | 0.31 ± 0.03 | 0.74 ± 0.03 | **0.61 ± 0.03** |
| MLS | AMRPlusPlus | 1 | 0 | 0.89 ± 0.02 | 0.50 ± 0.03 |
| | DeepARG | 0.87 ± 0.02 | 0.27 ± 0.03 | 0.85 ± 0.02 | 0.57 ± 0.03 |
| | Meta-MARC | 0.46 ± 0.03 | 0.33 ± 0.03 | 0.57 ± 0.03 | 0.40 ± 0.03 |
| | KARGA | 1 | 0.67 ± 0.03 | 0.96 ± 0.01 | 0.83 ± 0.02 |
| | **ResFinder** | 1 | 0.86 ± 0.02 | 0.98 ± 0.01 | **0.93 ± 0.02** |
| Phenicol | AMRPlusPlus | 0.93 ± 0.01 | 0.72 ± 0.03 | 0.88 ± 0.02 | 0.82 ± 0.02 |
| | DeepARG | 0.82 ± 0.02 | 0.29 ± 0.03 | 0.73 ± 0.02 | 0.55 ± 0.03 |
| | Meta-MARC | 0.33 ± 0.03 | 0.71 ± 0.03 | 0.44 ± 0.03 | 0.52 ± 0.03 |
| | **KARGA** | 0.86 ± 0.02 | 0.85 ± 0.02 | 0.88 ± 0.02 | **0.85 ± 0.02** |
| | **ResFinder** | 0.88 ± 0.02 | 0.88 ± 0.02 | 0.90 ± 0.02 | **0.88 ± 0.02** |
| Tetracyclines | **AMRPlusPlus** | 0.90 ± 0.01 | 0.85 ± 0.02 | 0.93 ± 0.01 | **0.88 ± 0.02** |
| | DeepARG | 0.90 ± 0.02 | 0.44 ± 0.03 | 0.87 ± 0.02 | 0.67 ± 0.02 |
| | Meta-MARC | 0.99 ± 0.01 | 0.45 ± 0.03 | 0.92 ± 0.01 | 0.72 ± 0.02 |
| | KARGA | 0.71 ± 0.02 | 0.90 ± 0.02 | 0.81 ± 0.02 | 0.80 ± 0.02 |
| | **ResFinder** | 0.94 ± 0.01 | 0.81 ± 0.02 | 0.94 ± 0.01 | **0.87 ± 0.02** |
| Diaminopyrimidines<br>(Trimethoprim) | **AMRPlusPlus** | 0.91 ± 0.02 | 0.91 ± 0.02 | 0.95 ± 0.02 | **0.91 ± 0.02** |
| | DeepARG | 0 | 1 | NA | 0.50 ± 0.04 |
| | Meta-MARC | 0.91 ± 0.02 | 0.09 ± 0.02 | 0.93 ± 0.02 | 0.50 ± 0.04 |
| | **KARGA** | 0.90 ± 0.02 | 0.91 ± 0.02 | 0.94 ± 0.02 | **0.90 ± 0.02** |
| | **ResFinder** | 0.91 ± 0.02 | 0.91 ± 0.02 | 0.95 ± 0.02 | **0.91 ± 0.02** |

Next, we evaluated the prediction concordance between each algorithm pair, which is illustrated in Figure 1. AMRPlusPlus, KARGA, and ResFinder cluster together, and exhibit moderate to high pairwise correlations, ranging between 0.62 and 0.72. Meta-MARC and DeepARG exhibit a lower mutual correlation (0.56), and are not well-correlated with the other methods, with a correlation ranging from 0.27 to 0.55.

**Figure 1:**
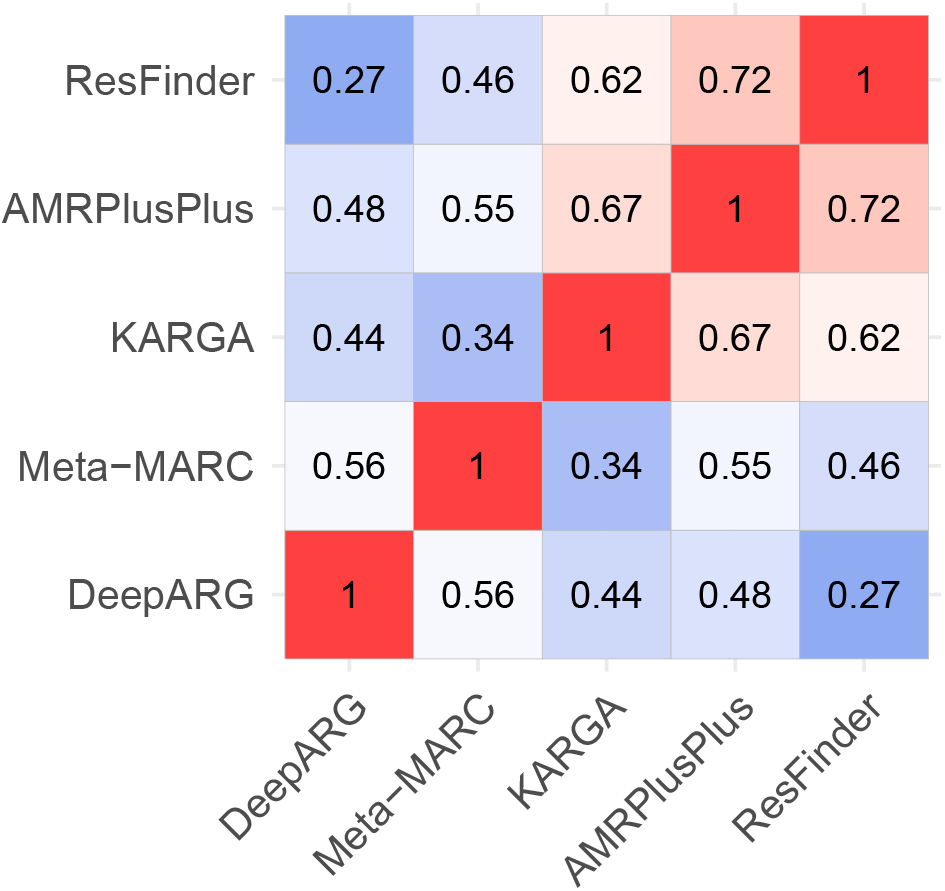
Heatmap of Pearson’s correlations between AMR classifiers’ predictions run on S1-S6 data sets (585 isolates).

To further investigate whether there are differences among algorithms even when they are concordant in overall resistance determination (e.g., if they identify two different AMR genes or mechanisms) we look at the most represented AMR groups by each algorithm, according to the MEGARes ontology (i.e., 42 groups for aminoglycosides, 221 for betalactamases, 17 for fluoroquinolones, 68 for glycopeptides, 83 for MLS, 18 for phenicols, 7 for sulfonamides, 62 for tetracyclines, and 11 for diaminopyrimidines). Since ResFinder and DeepARG have different categorizations, only AMRPlusPlus, KARGA, and Meta-MARC can be compared. Note that the group level is the finest AMR functional granularity that can be reached with these algorithms (AMRPlusPlus and KARGA can give gene predictions, but Meta-MARC cannot). Results are shown in Figure 2. Interestingly, even algorithms with higher correlation can identify very different groups at the base of the predicted resistance. For instance, in the fluoroquinolone class, KARGA finds primarily PATB and PATA, while AMRPlusPlus finds GYRA and PARC; in sulfonamides, FOLP is strongly predicted by both AMRPlusPlus and Meta-MARC, but not KARGA; on the other hand, only KARGA predicts SULI in a large number of isolates. Some of the differences may be explained by the fact that AMRPlusPlus and Meta-MARC are able to flag chromosomal genes that induce antibiotic resistance through point mutations, e.g., the GYRA topoisomerase enzyme subunit, while KARGA does not (because they need secondary confirmation of the point mutations). Another reason is that the presence of more than one gene could be needed for resistance, and the algorithms might identify one or the other, but not both. Finally, some genes could be very similar, leading to uncertainty in the identification and different frequencies in gene findings by different algorithms. Of note, Meta-MARC clusters very similar genes, and only one representative is used.

**Figure 2:**
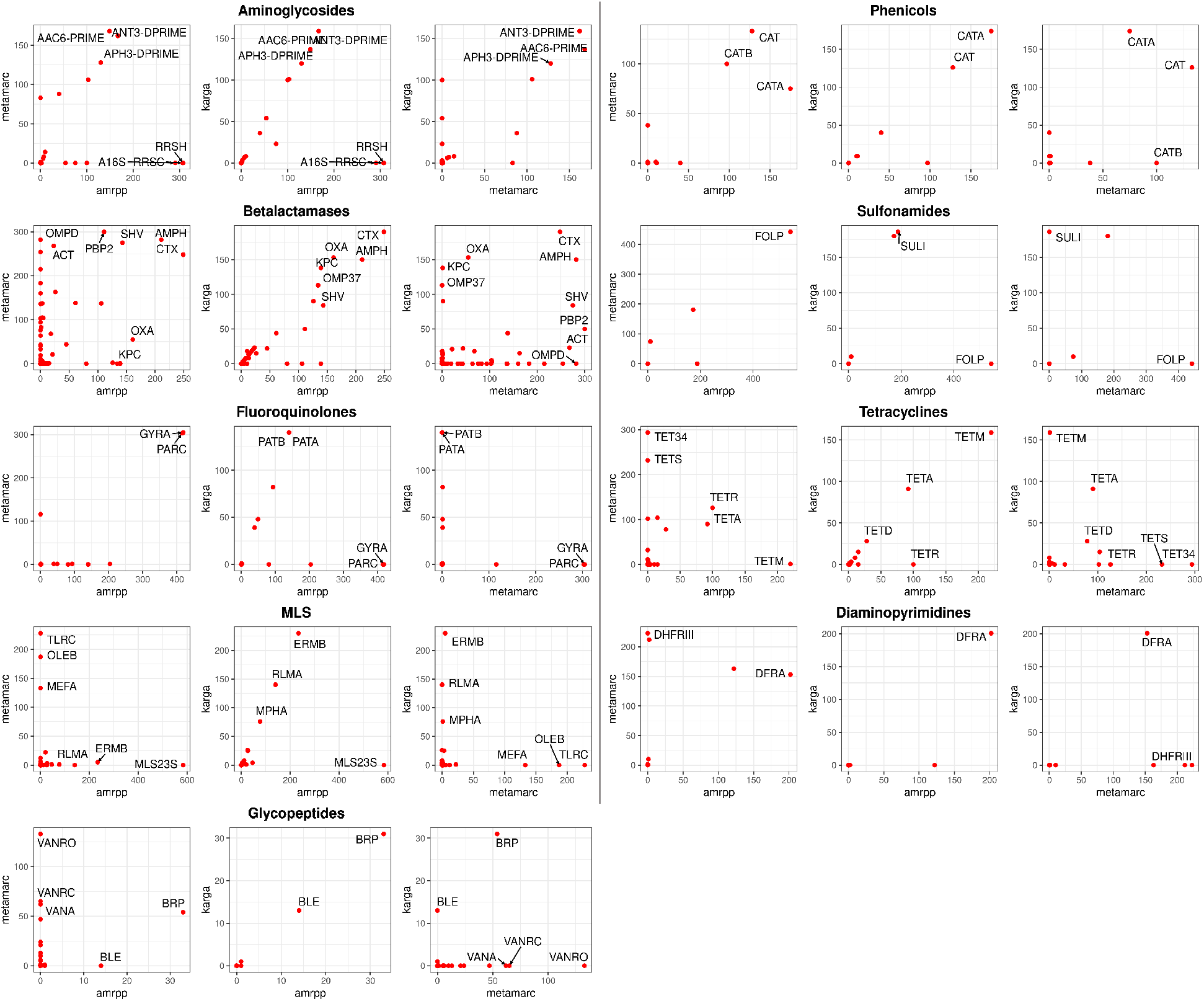
Per-class MEGARes AMR group prediction analysis on AMRPlusPlus, Meta-MARC, and KARGA. We measured the number of isolates predicted as positive for each group (regardless of the phenotype label), per class, indicated on the X and Y axes of the panels. All 585 isolates for each class are compared. We report the names of the top groups depending on the per-class group numerosity, as follows: top five per method if there are more than 99 groups in the class; top three if 99 to 25; top two if 24-15; and top one for less than 15 groups in the class. Of note, isolates without an associated glycopeptide AMR profile, are found resistant to the glycopeptide class (more than a hundred by Meta-MARC and more than thirty by KARGA and ResFinder).

#### Subgroup analysis based on isolation source

Our data set included 36 clinical isolation sources, namely Aspirate (1), Blood (10), Blood in broth (2), Built-environment (2), Conjunctiva (2), Generic patient sample (73), Deep wound swab (1), Ear swab (1), Groin (1), Gut (47), High vaginal swab (3), Hospital (9), Human-associated habitat (24), Low vaginal swab (2), Middle ear (45), Missing (3), Nasopharynx (38), No Label (24), Nose (7), Parotid gland (1), Perirectal Swab (4), Peritoneal Dialysis Fluid (1), Pus (11), R knee pus (1), Sink Aerator (1), Sink Drain (1), Sinus (2), Sputum (35), Throat swab (108), Throat/Groin (1), Urethral swab (1), Urine (19), Vaginal swab (2), Vulval swab (2), Hospital wastewater (98), and Wound swab (2).

To perform a subgroup analysis based on the isolation source, we analyzed the AMR predictions for the combinations of antibiotic class and isolation source presenting a total of at least 25 resistant and susceptible isolates. We retrieved six eligible subset combinations involving four AMR classes (aminoglycosides, fluoroquinolones, phenicols, and tetracyclines) and three isolation sources (hospital wastewater, generic patient sample, and throat swab). The combinations included aminoglycosides and hospital wastewater, fluoroquinolones and hospital wastewater, fluoroquinolones and generic patient sample, phenicols and generic patient sample, tetracyclines and throat swab, and tetracyclines and generic patient sample. The stratified BalAcc results are consistent with the aggregated ones reported in Table 4 for the majority of the combinations (KARGA produced top performance in four out of the six combinations, while ResFinder and AMRPlusPlus produced top performances in two and one out of the six combinations, respectively). However, a few specific combinations of AMR class and isolation source did heavily increase or decrease the quality of the predictions. As we mention in the Discussion, ad-hoc AMR data collections are needed to properly assess and train AMR detection algorithms so that they can be more resilient to the data collection context (including the isolation source). Detailed results for the different combinations of isolation source and antibiotic class are reported in Supplementary Table S5a-b, and in the Supplementary Methods.

#### Computational determination of AMR, prediction performance, time of execution, and agreement among algorithms

As described in the Introduction, timely execution is a crucial factor for introducing routine employment of computational AMR prediction tools. We assessed the speed of execution by first generating a “speed test” subset by subsampling ten isolates from each data set we considered in this study (10% of the whole data set, for a total of 60 compressed FASTQ pairs with mean size 357± 208MB). We then calculated the runtime for each algorithm on this subset. We ran the algorithms on 4 Intel(R) Xeon(R) CPUs E5-2698 v3 (2.30GHz), with 96GB of RAM, on the University of Florida HiPerGator 3 computing cluster. For each method, we measured the overall median (IQR) execution time in seconds, finding ResFinder to be the fastest with 105.50 seconds (80.75, 169.50), followed by KARGA with 190 (126.50, 263.20), DeepARG with 454 (197.20, 680), AMRPlusPlus with 920.50 (558.20, 1335.50), and Meta-MARC with 24,614 (14,466, 33,771). We also analyzed the results stratified by file size, considering inputs as compressed FASTQ pairs with sizes <0.50 GB (47 pairs), 0.50-1 GB (11 pairs), and >1 GB (2 pairs). We observe the stratified speed-of-execution rankings to be consistent with the unstratified results. We report the measured times in Supplementary Table S6.

## 3 Discussion

### Algorithm performance

Overall, all the methods except KARGA had a high number of false positive predictions, i.e., predicting the positive (resistant) class on negative (susceptible) isolates. In some specific cases, *all*, or almost all, the isolates for a class were classified as positive, i.e., betalactamases for ResFinder; aminoglycosides and fluoroquinolones for AMRPlusPlus, Meta-MARC, and DeepARG; and MLS for AMRPlusPlus.

We note that the classification results of DeepARG are given per read, i.e., the AMR class label is not predicted at the isolate level. Therefore, results must be aggregated, and a threshold applied. To better understand how this affected our results, in a sensitivity analysis, we considered two additional approaches: (a) using a higher threshold than the default probability (0.80) of DeepARG; and (b) requiring a minimum number of reads to support AMR determination at the isolate level. A threshold of up to 0.99 did not change the main results, i.e., in cases of positive classification, there was always at least one read with a predicted probability of 0.99 or more. For the second strategy, to label the whole isolate as resistant, we evaluated a grid of thresholds with a minimum number of resistant reads, i.e., 10, 100, 1000, and 10000. However, none of the values improved results consistently for all classes. For example, in the case of aminoglycosides, a threshold of 1000 led to the best BalAcc (0.53 versus the default of 0.50); the best thresholds for the betalactamase class were 1 and 10, all other values degraded the performance. In general, threshold optimization could lead to performance improvement, but we must note that: (a) there is not a one-size-fits-all solution, and the threshold must be optimized per AMR class; (b) this post-hoc adjustment may lead to overfitting, as the optimal threshold should be part of the training phase, and testing should be done on unseen data; (c) even optimizing the threshold, the results do not reach the performance of the top three algorithms in this study. Detailed results of the threshold-dependent predictions are provided in Supplementary Table S7. As a last note for DeepARG, the radical difference in prediction for diaminopyrimidines (trimethoprim), when compared to other algorithms, could be due to absence of specific trimethoprim sequences that are instead present in MEGARes.

We also verified whether tweaking the coverage parameter could improve the results on two problematic AMR classes, namely aminoglycosides and fluoroquinolones, where even the top two algorithms KARGA and ResFinder performed poorly due to low specificity. We therefore tested different values of the coverage parameter, to reduce the false negative rate. If we express coverage as a fraction, the default values are 0.60 for ResFinder and 0.80 for KARGA. We set a grid search by increasing the tools’ default values by 0.10, and re-ran the methods over our data set. We found that the optimization increased Specificity at the expense of Sensitivity, and the overall BalAcc had a modest increase at best. In detail, for aminoglycosides, KARGA reached a BalAcc of 0.68 (default: 0.66) when coverage was set to 0.90, while ResFinder achieved a similar BalAcc increase up to 0.66 (default: 0.64) when coverage was set to 1. For fluoroquinolones, increasing coverage in KARGA to 1 had only a negative effect on BalAcc: Spec increased to 0.71 (default 0.55), but Sens dropped to 0.32 (default 0.60); while with a coverage of 0.90, Resfinder increased BalAcc from the default 0.61 to 0.62. It must be noted that these modest increases are subject to the aforementioned limitations (a-c). Detailed results of the coverage threshold-dependent predictions are provided in Supplementary Table S8.

In this study we compared the performances of high-throughput algorithms that work directly with short read sequencing data. Of note, there is another class of algorithms which take whole bacterial genomes or assembled contigs as input, either as nucleotides or translated protein, such as VAMPr [17] and PARMAP [18], respectively. In principle, these algorithms could be used with WGS high-throughput sequencing by performing genome assembly, although they would not work for metagenomics sequencing experiments, which could be performed in certain clinical settings. To obtain genome assemblies from short reads followed by protein translation, several extra steps are required. These steps often need to be done independently by the user, e.g., the software code/pipelines provided by VAMPr and PARMAP do not include them. Note that genome assembly does not involve the application of just one program, e.g., de novo or reference assembly, but involves pre-assembly quality control and error correction, parameter optimization of the assembly itself, and scaffolding/finishing after assembly. The extra steps may heavily influence the downstream results for an AMR prediction algorithm. In other words, a high-quality assembly software pipeline could boost the performance of VAMPr and PARMAP, while a low-quality assembly software could have the opposite effect. For proper algorithm comparison, and to avoid providing unfair advantage, we ran all the pipelines with the developers’ recommended settings. Adding assembly and protein translation phases would add a large, unavoidable user-choice component to the analysis presented in this study. For these reasons we consider VAMPr, PARMAP, and other algorithms based on full genome inputs to be out of the scope of the present review. Even if these algorithms are limited to well-curated full genomes, and selected combinations of pathogens and antibiotics, they are competitive: VAMPr showed very high accuracy (averaging 0.91 in validation sets), and PARMAP provided both AUC and Recall higher than 0.80. We note also that Meta-MARC, included in this study as it can process both assembled and unassembled data, retrieved more AMR genes in the validation sets when assembled contigs were used instead of short reads [11].

In summary, we observed that no one algorithm provides outstanding results in all of the considered AMR classes in terms of balanced accuracy, and the same holds for the *f*_1_ measure. Yet on average they all perform well. Although encouraging, these results are still far from a near-perfect classification that would be needed to include them in clinical practice as substitutes to phenotypic AST.

### Distribution shift and bias

All high-throughput AMR prediction algorithms, based on sequence alignment, *k*-mers, or machine learning, are trained on an established ground truth. Training data sets include well-characterized, laboratory-confirmed AMR genes, such as MEGARes[9] or CARD[5], or collections of whole genomes with associated AST phenotypes, as in PATRIC[7]. They are the result of a coordinated effort by the scientific community to collect and standardize AMR knowledge, but can be subject to several types of bias, sampling in primis[10, 20]. For example, a genus or species can be over-represented; distribution of samples over time are not uniform; and there is substantial geographical bias (by country or samples within localized outbreaks). Species distribution shifts can be common even within related clinical settings, e.g. in infections whose source/colonization origin is different or when treatment protocols change over time. Evidence of bias across genus, temporal, and geographical representation was found in PATRIC, and it was shown that such bias affects discriminative ability of AMR machine learning predictions by 5% [20]. For most of the AMR classes in our data set we found strong geographical and per-genus distribution shifts. We consider this finding a further indication that these sources of bias should be accounted for when collecting data used as a reference or training set for AMR prediction algorithms.

### Problems with ground truth

Similar to standardized AST methods, there is a need to develop standardized AMR bioinformatics protocols and evaluation benchmarks, so that results from different studies can be put together and used for training and validation. So far, the Food and Drug Administration does not provide standardized regulatory guidelines for metagenomic diagnostic tests[3]. A second major obstacle to comparing the performances of different AMR classification tools is that they can be based on different AMR ontologies –albeit an output match is possible for most antibiotic classes at different granularity levels. We must note that CARD and MEGARes ontologies are designed to organize genes and resistance mechanisms, thus they are well suited for studies on metagenomics and ecology, but less for clinical studies where single species are cultured and tested against antibiotics. Although there are other off-the-shelf tools that predict directly AST from genotypes, they need assembled and finished genomes, which is a non-trivial (and time-consuming) step in the analytics pipeline, while all the methods used here accept raw short read data.

The high-throughput AMR prediction algorithms we examined provide mostly a per-AMR-class prediction, with a few exceptions, e.g., MEGARes-based output below the class level, such as the specific penicillin binding protein mechanism. However, the isolates we consider have been tested only against some specific antibiotics within a class. In the case of resistant isolates, this is not necessarily a problem. For example, if an isolate is resistant to ampicillin, since it is resistant to at least one antibiotic of the betalactamase class, we can consider it resistant to the whole class and the algorithmic recommendation would be not to use betalactamases to address this particular infection. This assumption does not hold, however, for susceptibility. If another isolate is, for example, susceptible to ampicillin, we cannot label it with certainty as susceptible to all betalactamases –in order to do so, we should in fact test it extensively against a large fraction (ideally, all) of the betalactamase class antibiotics. This poses another a priori problem with the ground truth, namely the fact that susceptible instances might be falsely labeled as they were not tested against all antibiotics of each specific class. It is however unrealistic to expect from observational studies, such as the ones we used to assemble the data set used for this work, to explicitly test resistance against an exhaustive range of antibiotic classes and molecules. Even large genome collections show how most of the species versus antibiotic combinations have few to no records[7, 10]. Another problem in assessing the correctness of the predictions is that the AMR framework description can include labels that are not informative enough to infer antibiotic resistance at the class level. This is the case, for example, of MEGARes multi-compound or multi-drug classes, or the DeepARG ‘unknown’ category. Such annotations are not suitable for a direct integration within clinical practice, as they do not provide precise indications to which AMR classes the sample or isolate is resistant to.

## 4 Conclusions

Current computational tools for AMR characterization from high-throughput sequencing data show promising results, but do not appear ready for application in clinical settings. There are obstacles that go beyond algorithmic development and relate to bias in data as well as issues with ground truth and ontology, hampering algorithm development and benchmarking. Therefore, we make a call to develop prospective studies designed with the specific intent of training and validating computational AMR tools, i.e., with explicit determination of possible sources of bias, comprehensive characterization of AST profiles, and linkage with at least one of the current ontologies. These data sets would then constitute the gold standard ground truth to build upon AMR detection algorithms.

## 5 Key points

- Current computational tools for AMR characterization from high throughput sequencing data show promising results but do not seem ready for application in clinical settings.
- Each method provides a wide range of performances over the considered AMR classes; no method seems to provide the best results in all the classes.
- The major roadblocks towards a realistic bench-to-bedside implementation are:
  - The presence of bias, such as species, geographical, and temporal biases in the reference/training data sets
  - The need to establish a per AMR-class ground truth by testing an extensive portion of the antibiotics (ideally, all of them) for each specific antibiotic class
  - Absence of standardization in both AST and AMR ontologies

## Acknowledgment

The authors want to thank Marco Oliva, Ilya B. Slizovskiy, and Daniele N. Cudin for the productive and deep conversations.

